# ANDOR and beyond: dynamically switchable logic gates as modules for flexible information processing

**DOI:** 10.1101/2021.08.02.454761

**Authors:** Mohammadreza Bahadorian, Carl D. Modes

## Abstract

Understanding how complex (bio-)chemical pathways and regulatory networks may be capable of processing information in efficient, flexible, and robust ways is a key question with implications touching fields across biology, systems biology, biochemistry, synthetic biology, dynamical systems theory, and network science. Considerable effort has been focused on the identification and characterization of structural motifs in these signaling networks, and companion efforts have instead sought to cast their operation as controlled by dynamical modules that appear out of dynamical correlations during information processing. While both these approaches have been successful in many examples of biological information processing, cases in which the signaling or regulatory network exhibits multi-functionality or context dependence remain problematic. We here propose a small set of higher-order effective modules that simultaneously incorporate both network structure and the attendant dynamical landscape. In so doing, we render effective computational units that can perform different logical operations based purely on the basin of attraction in which the network dynamics resides or is steered to. These dynamically switchable biochemical logic gates require fewer chemical components or gene products overall than their traditional analogs where static, separate gates are used for each desired function. We demonstrate the applicability and limits of these flexible gates by determining a robust range of parameters over which they correctly operate and further characterize the resilience of their function against intrinsic noise of the constituent reactions using the theory of large deviations. We also show the capability of this framework for general computations by designing a binary adder/subtractor circuit composed of only six components.

## 1 Introduction

Biology is absolutely filled to the brim with information processing systems of all types, at all levels of complexity, and across all relevant length and time scales^1^. At the level of neuronal information processing, pack animals navigate complex social decisions requiring integration of vast amounts of information about other members of the cohort^2,3^; meanwhile predators must make rapid, complicated decisions during their hunt for food while potential prey must navigate similar complexities to avoid being predated upon^4,5^. Similarly, in the realm of biochemical information processing, individual cells respond to small changes in the concentration of signaling molecules in non-trivial, non-linear ways via the biochemical signaling networks^6^. And the expression of the genes themselves are controlled by vast gene regulatory networks which respond to and incorporate significant amounts of contextual information^7,8^. Indeed, the knife’s edge of natural selection is so sharp that seemingly any trait or pathway or regulatory task whose effectiveness or efficiency could have been improved by incorporating non-trivial information processing has been. Unsurprisingly, therefore, the study of information processing in biology has a rich and varied history from neuronal systems^1^, and the establishment of layered neural network architectures that led to deep learning techniques to the exploration and enumeration of a significant fraction of the biomolecular players that participate in intra- and extracellular regulatory pathways^9^. The identification of prevalent circuit motifs in both networks of neuronal connections and in the regulatory networks has been a particularly important advance and major interest and effort continues in this vein today.

The overwhelming majority of these efforts, however, conceptualize information processing in biological systems as occurring over static, established circuits; one circuit performs one information processing task. In order to alter the information processing task performed by the circuit, the circuit architecture itself (the network topology or the edge strengths) or the control parameters must therefore be changed^10^. But this is not the only possibility. In the world of silicon-based information processing, for example, field programmable gate arrays, in which the memory bits in the control layer set the connections between the logic gates in the circuit layer, are not constrained by static circuit topologies^11^. Recently, neuroscience as well has seen renewed attention to the idea of flexible or dynamically-sourced information processing, in order to meet the challenge of mounting experimental results indicating attention or context-based task switching on timescales incompatible with plastic network adaptation^12^.

And what of signaling and regulatory networks? Here the supremacy of the one circuit, one task orthodoxy remains mostly unchallenged, with only relatively few studies considering the role of the underlying dynamics^13–15^. We instead here propose a new class of module decomposition of signaling and regulatory networks that incorporates both topological structure and the dynamical landscape playing out over that structure. These modules can be thought of as *dynamically switchable logic gates*, where the basin of attraction of the dynamics determines the information processing task that the network performs. These tasks are further mappable directly to the classical logic gates that comprise an effective basis for all of information processing.

The fact that in these systems the function may be affected by the dynamics could have many benefits, such as enabling switching among different tasks at a higher speed^16,17^. Moreover, it circumvents the need for rerouting signal to a specific unit which is particularly difficult in many biological cases, where a signal is the abundance of a biochemical species available everywhere. Additionally, since such complex systems generally have several functions, being able to perform many of them with a given subsystem is crucial. Especially, when there are limited resources as in most practical and natural examples. Therefore, in this study we propose a framework for studying context-dependent information processing which allows different procedures (e.g. logical functions) by the same unit on a given input.

Studying decision making in a given system typically benefits from investigating the building blocks of information processing, i.e. *logic gates*, in that framework. For example, discussions about quantum information processing devices have been based on their constitutes: quantum logic gates^18^. Similarly, many studies have been focusing on designs and properties of logic gates which can be potentially implemented by biological systems^19–21^.

While traditional logic gates make a useful toy model for studying some aspects of information processing devices, they are unable to constitute a multifunctional circuit. In other words, dynamically switchable logic gates are required that can dynamically switch between different functions based on the demand. These circuits are particularly important for studying the cases in which the device is performing multiple functions and the switching action occurs due to the dynamics of the system^22^. Changing the function this way, in examples such as neural networks^23^ or ecosystems in shallow lakes^24^, enables the system to switch much faster compared to changing the structure (e.g. due to training). It should be noted that although some of the currently available gate designs have the capability of multifunctional operation, e.g. by increasing the output’s threshold, a gate can operate as AND and OR gates^25^, a systematic study of such multifunctionality and their costs and benefits is missing. Moreover, the switching in these designs are dictated externally and directly to the output species by changing its inhibitor concentration. This is equivalent to altering the circuit’s structure, and it is not a result of the underlying dynamics of the system. Therefore, in this article, we introduce some examples of *dynamically* switchable logic gates, show their applicability by constructing a binary adder/subtractor, and then, discuss advantages and disadvantages of multifunctionality based on these circuits.

## 2 Results

### 2.1 Generic design of dynamically switchable logic gates

Let us begin with a generic configuration of a dynamically switchable logic gate, as depicted in Fig. 1, similar to the proposed circuit in Ref.^22^ in the context of neural networks and their attendant dynamics. In our case, each element of the circuit (shown by an orange circle) represents a biochemical species that can be a gene product or any other regulatory component of the cells with switch-like behavior as discussed in Sec. 4.1. In this configuration, an intermediate layer (in the green box) receives the signal from two inputs (e.g. upstream genes), and sends a signal to the output (e.g. a downstream gene) according to its state variables. In this setup, the positive signal (promotion) from the two components in the intermediate layer is necessary for activating the output. It should be noted that in this Fig. an the following ones the links have unit weight unless it stated otherwise in the diagram. Besides, throughout this article for simplicity, we consider the state of the left layer (inputs to the intermediate) to be static, independent from each other and acquiring either zero or one corresponding to OFF and ON, respectively. We here need only focus on the dynamics and fluctuations of the intermediate layer.

**Figure 1.**
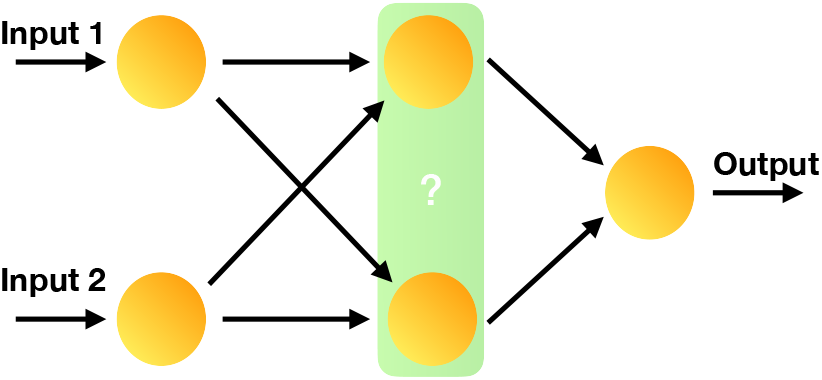
The structure of the proposed generic dynamically switchable logic gate.

### 2.2 The ANDOR gate: structure and dynamics

The simplest case is when the logic gate can switch between “AND” and “OR” functions here called ANDOR. According to the truth tables of these gates, the output should be OFF regardless of the gate type when there is no input. Similarly, for both gate types, the output is ON when both inputs are ON, and the only difference between these two gates is in their response to the intermediate signal level, i.e. when one of the inputs is ON and the other is OFF. In this case, the OR gate should result in ON output while the AND gate’s output should be OFF. Therefore, replacing the green box in Fig. 1 with a circuit which has bistability for intermediate input signals enables the gate to perform both OR and AND tasks. This can be achieved by replacing the green box of Fig. 1 with a bistable motif, as shown in Fig. 2a. Note that here, *s* is the sum of the two inputs to the intermediate layer (i.e. *s ≡*Input 1 + Input 2) which can only be zero, one or two and which we assume to be static. However, the concentration of the two genes (i.e. *x*_1_ and *x*_2_) in the intermediate layer can have any positive real values and their dynamics are given by

**Figure 2.**
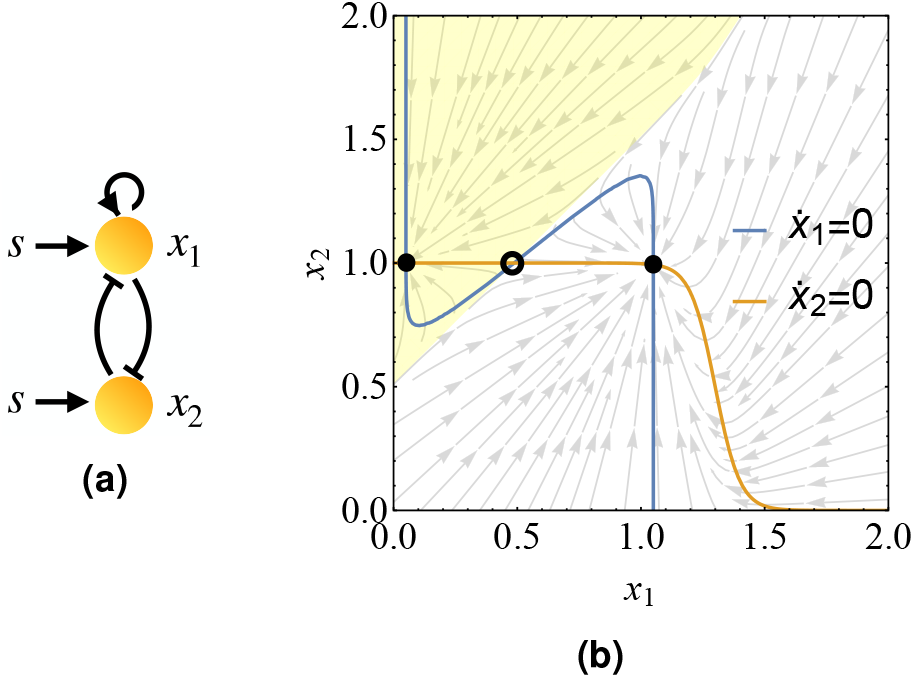
**(a)** Topology of a bistable motif in which one of the stable fixed points corresponds to both genes being expressed. Here, *s ≡* Input 1 + Input 2. **(b)** The phase portrait of the bistable motif shown in (a) with intermediate signal *s* = 1. The black filled and hollow circles show the stable fixed points and saddle point, respectively. Moreover, the shaded area shows the basin of attraction of the left fixed point.

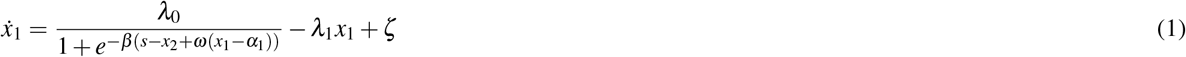

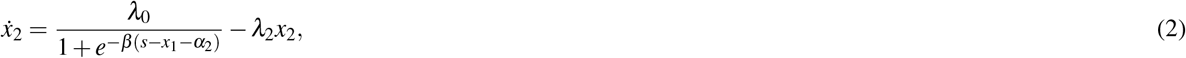

where *ω* controls the slope of the separatrix, and *λ*_1_ and *λ*_2_ are the degradation rates for *x*_1_ and *x*_2_, respectively. *λ*_0_ is the production rate of both genes when they receive a strong enough input. *α*_1_ and *α*_2_ are the activation thresholds for *x*_1_ and *x*_2_ which in our case are equal to 0.5 and 0.3, respectively. Finally, *ζ ≪* 1 is needed for numerical stability. See Sec. 4.1 for more details and the basic idea of this type of dynamics.

This motif satisfies the required conditions mentioned for no input and two inputs with a proper set of parameters. In order to check the bistablity region, required for the intermediate input level (i.e. *s* = 1), one can simply check the phase portrait. In this type of plots, a typical trajectory for every small neighborhood of the phase space is represented by an arrow whose size is proportional to the distance covered by the system in a given time interval. In Fig. 2b, we have the phase portrait of the ANDOR gate along with the curves that show 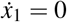 and 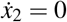. These two curves cross each other three times producing three fixed points. Two of them are stable fixed points (shown by filled black circles) separated by another one which is a saddle point (shown by a hollow circle). The stable fixed point at the center corresponds to the situation in which both genes are expressed and available for the downstream genes while the other stable fixed point corresponds the situation in which only *x*_2_ is expressed, and it represses the expression of *x*_1_. Therefore, replacing the green box in the generic configuration in Fig. 1 with this motif enables the system to perform both AND and OR functions, because when only one input is ON, the output can be ON or OFF depending on the fixed point that the intermediate genes *x*_1_ and *x*_2_ reach. If they approach the fixed point in the center, the output will be ON and it means that one input is enough for making output ON, i.e. the system works as an OR gate. On the other hand, if the genes reach the left stable fixed point, the total signal for expression of the output gene is not enough and the whole circuit functions as an AND gate. Whether the ANDOR gate performs AND or OR function depends on the “context” of the decision and is discussed in the Sec. 2.2.1.

It should be noted that the ANDOR gate only requires five components while traditional static AND and OR gates together require six. Besides, an additional controller unit is needed to redirect the signal to the desired gate if function switching is not possible. Therefore, using the ANDOR gate can reduce the number of required components significantly.

#### 2.2.1 Logical function switching in the ANDOR gate

In this system, switching between these two functions is possible by simply transiting from one basin of attraction (shaded area or the rest of the phase space in Fig. 2b) to the other one. This transition can happen due to an additional and external signal to gene *x*_2_ which might be tissue specific and originate from the extracellular environment. By such a signal, the expression of *x*_2_ can be controlled and the system can be steered vertically to the desired basin of attraction. With such a strategy, cells would potentially be able to make different decisions depending on the tissue which they are part of, and the external signal that they receive. Moreover, the decision made (i.e. the approached fixed point) also depends on the initial content of expressed genes. This strategy could be employed in cell fate decision making processes where two daughter cells acquire different fates due to asymmetric distribution of cell contents during the division. For example, a transcription factor of *x*_2_ could be divided asymmetrically between two daughter cells, giving rise to two distinct decisions and cell fates. However, if the system is obliged to start from the origin of the phase space, the final state can be controlled by adjustment of the slope of the separatrix (the line which separates two basins of attraction). In this scenario, the function (i.e. AND or OR) that the system performs can be interpreted as a “ground state” of the system since it is the function that system performs naturally, without requiring injection of any additional external energy for steering the system.

#### 2.2.2 Characterization of the ANDOR gate

In order to understand the potential application of this multi-functionality, one needs to explore the phase diagrams of the system. Fig. 3a shows the different regions in the dimensionless parameter space of the *sβ* and 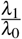 plane when 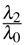 is set to 1. The blue area shows the parameter combinations which satisfy all conditions required for performing AND and OR functions. The yellow part shows the region in which the system at the central fixed point does not send enough signal to the output (i.e. *x*_1_ + *x*_2_ *<* 1.5) which means that the system can not perform the AND function. Finally, the red region is where the bistability does not exist. Similarly, we also determine the right combination of 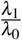 and 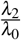, as shown in Fig. 3b for *sβ* = 20. Here, each color has the same meaning as as in Fig. 3a, and the gray color shows the region where degradation for both *x*_1_ and *x*_2_ is so low that even without input, their steady-state concentrations meet the output threshold and it turns ON. As one can see in these figures, there is a robust range of parameters for which this circuit behaves as a context-dependent logic gate switching between AND and OR.

**Figure 3.**
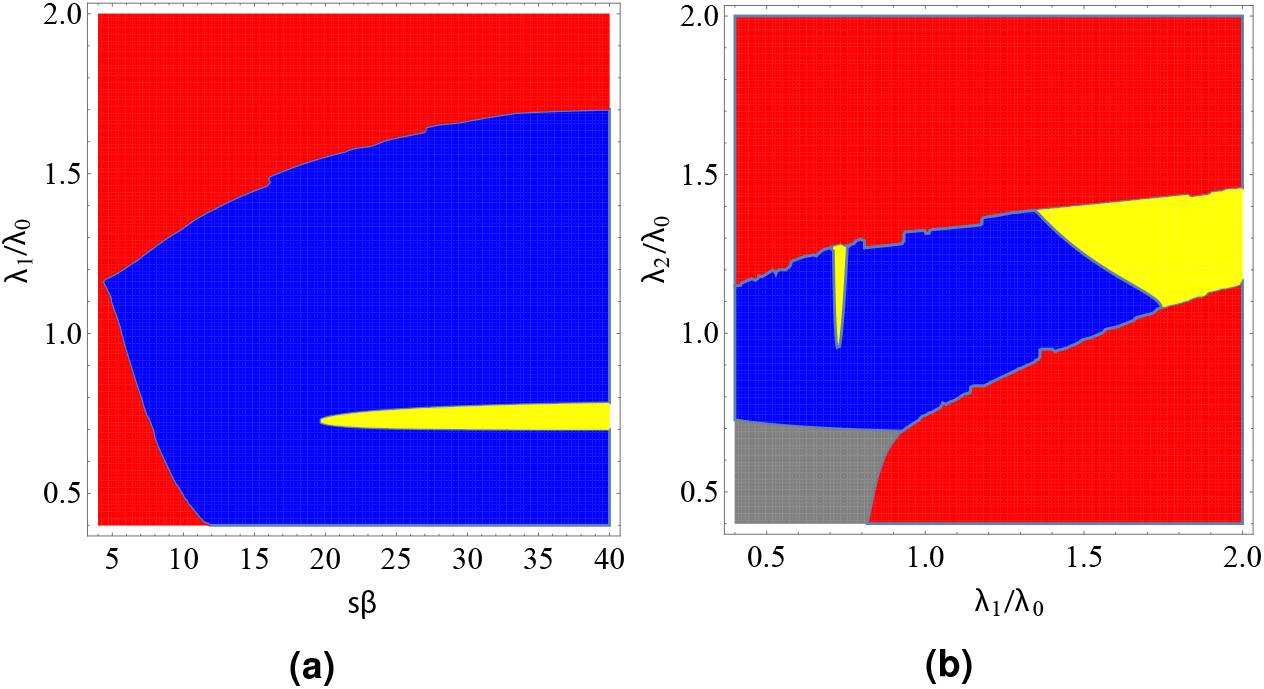
Phase diagram of the system described by Eqs. 1 and 2 in **(a)** *sβ* – *λ*_1_*/λ*_0_ and **(b)***λ*_1_*/λ*_0_– *λ*_2_*/λ*_0_ planes. Blue color shows the region in which system meets all requirements while red area shows where it does not have bistability. When the bistability exists, yellow shows where the sum of signals to the output does not meet its threshold, and gray shows where the output turns ON without any input (i.e. *S* = 0). Note that in both phase plots the parameter range for which the network operates as an ANDOR gate is significant.

#### 2.2.3 A MAP for the ANDOR gate: noise-induced transitions and robustness of the operations

Although the aforementioned bistability enables the system to perform two distinct functions with a wide set of parameters without requiring twice as many components, it also allows undesired noise-induced transitions (i.e. errors) that reduce the reliability of the decisions. In order to quantitatively study these transitions, one can minimize the Freidlin-Wentzell action to find the most probable path taken by the system for a transition as well as the probability of that path up to a normalization prefactor when the noise strength is small. See Sec. 4.2 for more details.

Due to random timings of the chemical reactions in biological systems, the concentrations of species follow a stochastic dynamics. In the case of well-mixed systems, one can use the chemical Langevin equation^26^ to fully describe the dynamics when fluctuations are sufficiently small (i.e. the reaction volume is large). For the ANDOR gate whose dynamics is described by Eq. 4, the action over the path *φ* can be written as

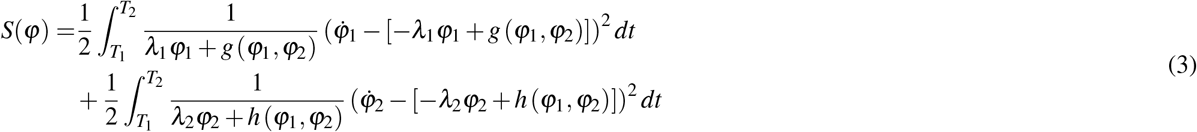

in which *φ*_1_ and *φ*_2_ are the coordinates of the path *φ* at any time *t ∈* [*T*_1_, *T*_2_]. *g* (*φ*_1_, *φ*_2_) is the expression (the first term in the r.h.s. of Eq. 1) of *x*_1_ and *h* (*φ*_1_, *φ*_2_) is the expression of *x*_2_ (the first term in the r.h.s. of Eq. 2). The probability of this path to be taken by the system is proportional to 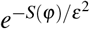 where *ε* is the noise strength and here equals to Ω^*−*1*/*2^ while Ω is the reaction volume. When Ω is large, all paths from a given point to another chosen one will have negligible probabilities compared to the one which minimizes the action in Eq. 3. This path, also known as the Minimum Action Path (MAP), determines the path with the highest probability for a given transition.

The MAP which connects the central fixed point to the left one for a typical set of parameters is shown in Fig. 4. The color on the path shows the gradient of the action at any given point. This gradient can be interpreted as the effective force applied by the noise for causing the movement along the MAP. As one can see in this figure, the MAP goes directly towards the separatrix in the opposite direction of stream lines. Then, it follows the streamlines close to the separatrix until it approaches the saddle point, and finally enters the other basin of attraction and follows those streamlines to the left fixed point. The observation that the MAP for the ANDOR gate crosses the separatrix close to the saddle point is consistent with the findings of other studies in different contexts^27^.

**Figure 4.**
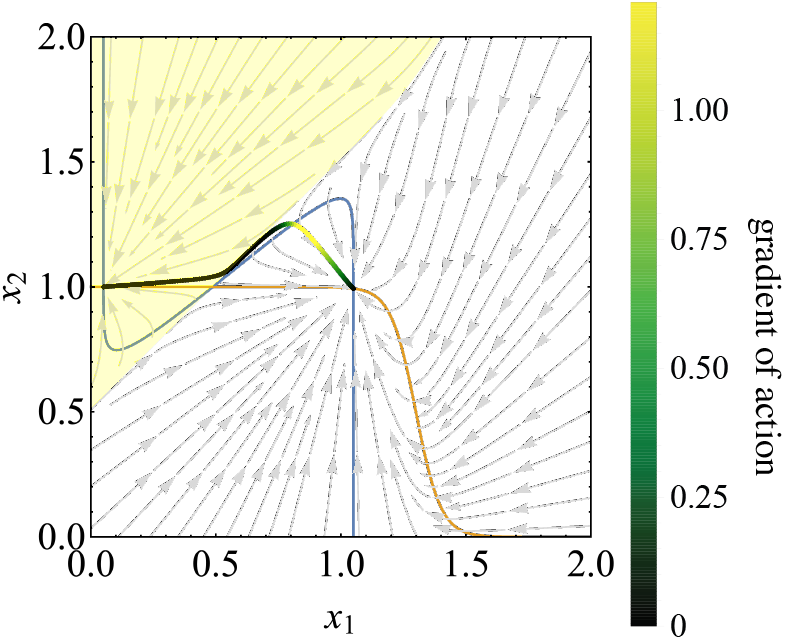
Phase portrait of the ANDOR gate shown in Fig. 2 along with the MAP which connects the central fixed point to the left one.

The rate of the noise-induced transitions is a proxy for the reliability of the decisions made by the system. Therefore, beside exploring whether the truth tables of the gates are satisfied, we also investigate this reliability (i.e. resilience against noise) of the system as the parameters change. Fig. 5a show the action vs. sharpness of transition *β* for the transitions from the left fixed point to the right one (in black circles) and vice versa (in red triangles). As one can see in this figure, as *β* increases, the action for both transitions increase, but for the L *→* R transition increases faster which results in an critical value at *sβ*^*∗*^ = 14.75. It should be noted that although higher values of action *S* mean higher reliability for the decisions, any difference between the actions for two transitions result in a bias in the steady state of probability distributions. The dependence of the action *S* (*φ*) on the slope of the sepratrix *ω*, degradation rate *λ*_1_ of *x*_1_, and that of *x*_2_ are shown in 5b, 5c, and 5d, respectively. The critical values at which the action for L *→* R transition equals R *→* L are *ω*^*∗*^ = 0.87, 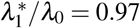 and 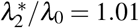. Note that in all these figures, unless a parameter is the variable of the plot we set them as following: *sβ* = 20 and *λ*_1_*/λ*_0_ = *λ*_2_*/λ*_0_ = *ω* = 1. Parameters consistent with one strongly stabilized basin at the expense of the other are attainable. On the other hand these parameters could be poised at criticality in order to equalize the transition probability and minimize the bias. Rapid switching would therefore be promoted in this scenario.

**Figure 5.**
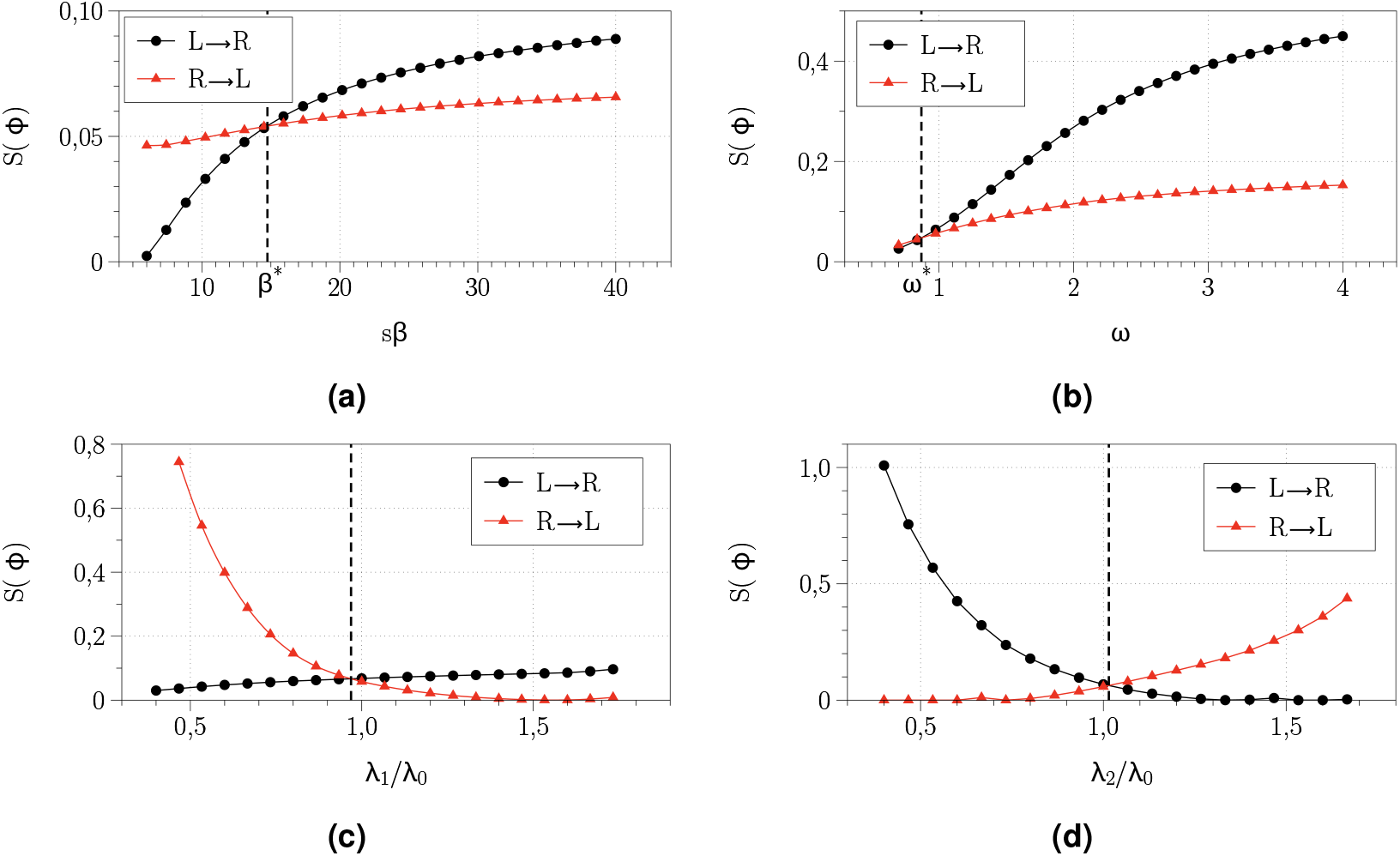
Dependence of minimum action for noise-induced transition **(A)** on the dimensionless sharpness of expression *sβ*, **(B)** on the slope of separatrix *ω*, **(C)** on the degradation rate *λ*_1_*/λ*_0_, and **(D)** degradation rate *λ*_2_*/λ*_0_. Black circles show the action for the transition from the right fixed point the left one and the red triangles show the action for the opposite transitions. For each dependency a critical value at which the cost of transiting L *→* R and R *→* L is balanced can be attained and is indicated by dashed line and have the following values: *sβ*^*∗*^ = 14.75, *ω*^*∗*^ = 0.87, 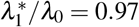 and 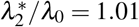.

### 2.3 The ANDOROFF and XORANDOROFF gates: Increasing the complexity and multifunctionality of the circuits

By increasing the complexity of the intermediate layer (green box in Fig. 1), one can design circuits which are able to switch among a greater number of distinct functions. As an example, consider the situation in which both genes in the toggle switch have a self-induction loop, as depicted in Fig. 6a. In this circuit, for the intermediate signal level (i.e. *s* = 1) there are three stable fixed points where two of them are identical to the ones in Fig. 2 while the third stable fixed point corresponds to a state in which *x*_1_ dominates and inhibits the expression of *x*_2_. Moreover, when *s* = 2 (i.e. both inputs are ON), one of the fixed points in which output is OFF will be preserved (as shown in Fig. 6b) if the self promotion loop of *x*_2_ has a relatively high strength, for example twice as strong as the other promoting links. The basin of attraction of this fixed point (shown in yellow) corresponds to the set of initial conditions from which the system cannot reach the central fixed point (neither with one input nor with two). Thus, this area represents the situations where the system acts as an OFF switch. This means that adding a new self loop and producing a new fixed point enables the system to switch among three different functions: AND, OR and OFF. We therefore name it an “ANDOROFF” gate.

**Figure 6.**
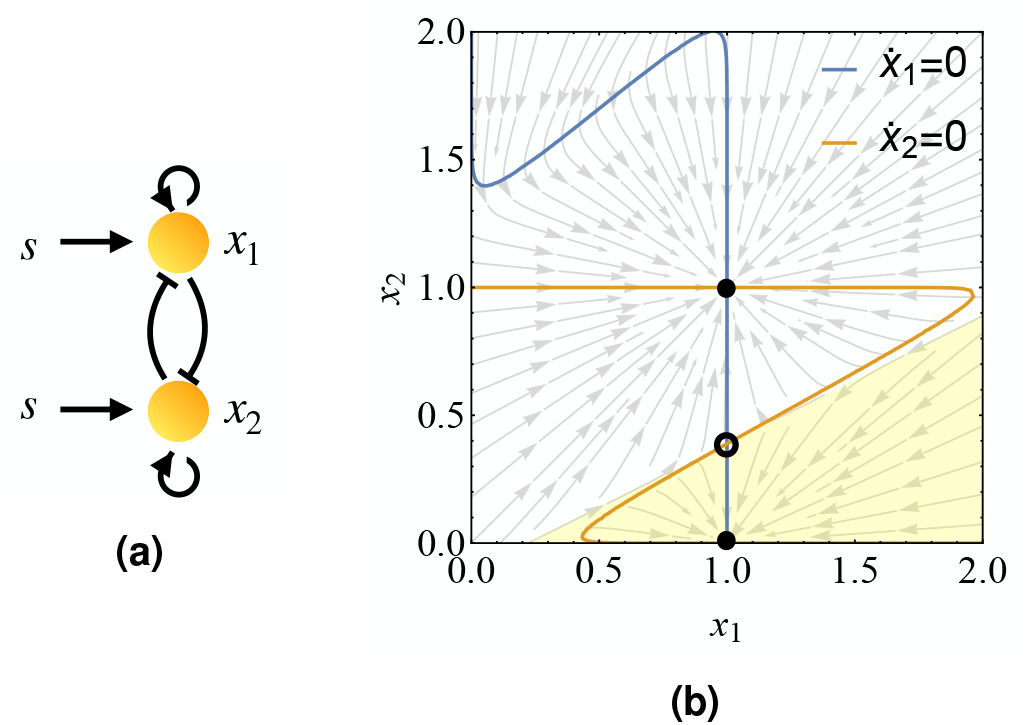
The ANDOROFF gate. **(a)** Topology of a multistable motif which has three stable fixed points for intermediate signal level and two for high signal level which enable it to perform AND, OR and OFF logical functions.**(b)** The phase portrait of the bistable motif shown in (a) with high signal *s* = 2. The black filled and hollow circles show the stable fixed points and saddle point, respectively. Moreover, the shaded area shows the basin of attraction of the lower fixed point.

Although increasing the number of fixed points can enable the system to perform more functions, it decreases the size of the basin of attraction of each fixed point. Accordingly, the reliability of each decision, defined as the robustness of each fixed point against uncertainty in the initial conditions, decreases. One way to overcome this limitation is by increasing the dimension of the phase-space by increasing the number of regulatory components at the intermediate layer. For example, the “XORANDOROFF” gate depicted in Fig. 7a has three genes in this layer which, with a proper set of parameters, result in four fixed points ((0, 0, 0), (1, 0, 0), (0, 1, 1) and (1, 1, 1)), in total. However, for each input signal (*s* = 0, 1 or 2), only two of them are available, and with the appropriate set of parameters, this circuit can switch between four functions: XOR (exclusive OR), AND, OR and OFF. In Fig. 7b, consider a system initially located at the blue region. For no input the system approaches the fixed point at which no gene is expressed (0, 0, 0) and output is OFF, and when one of the inputs is ON, the system, starting from this region, approaches the fixed point at which only *x*_2_ and *x*_3_ are expressed (0, 1, 1) and therefore, the signal for output is strong enough to make it ON. However, when both inputs are ON all three genes (*x*_1_, *x*_2_ and *x*_3_) will be expressed and expression of *x*_1_ inhibits the expression of output. Therefore, if the system is initially located in the blue region it functions as an XOR gate. Similarly, locating the system in other regions enables it to perform other functions and to make different decisions.

**Figure 7.**
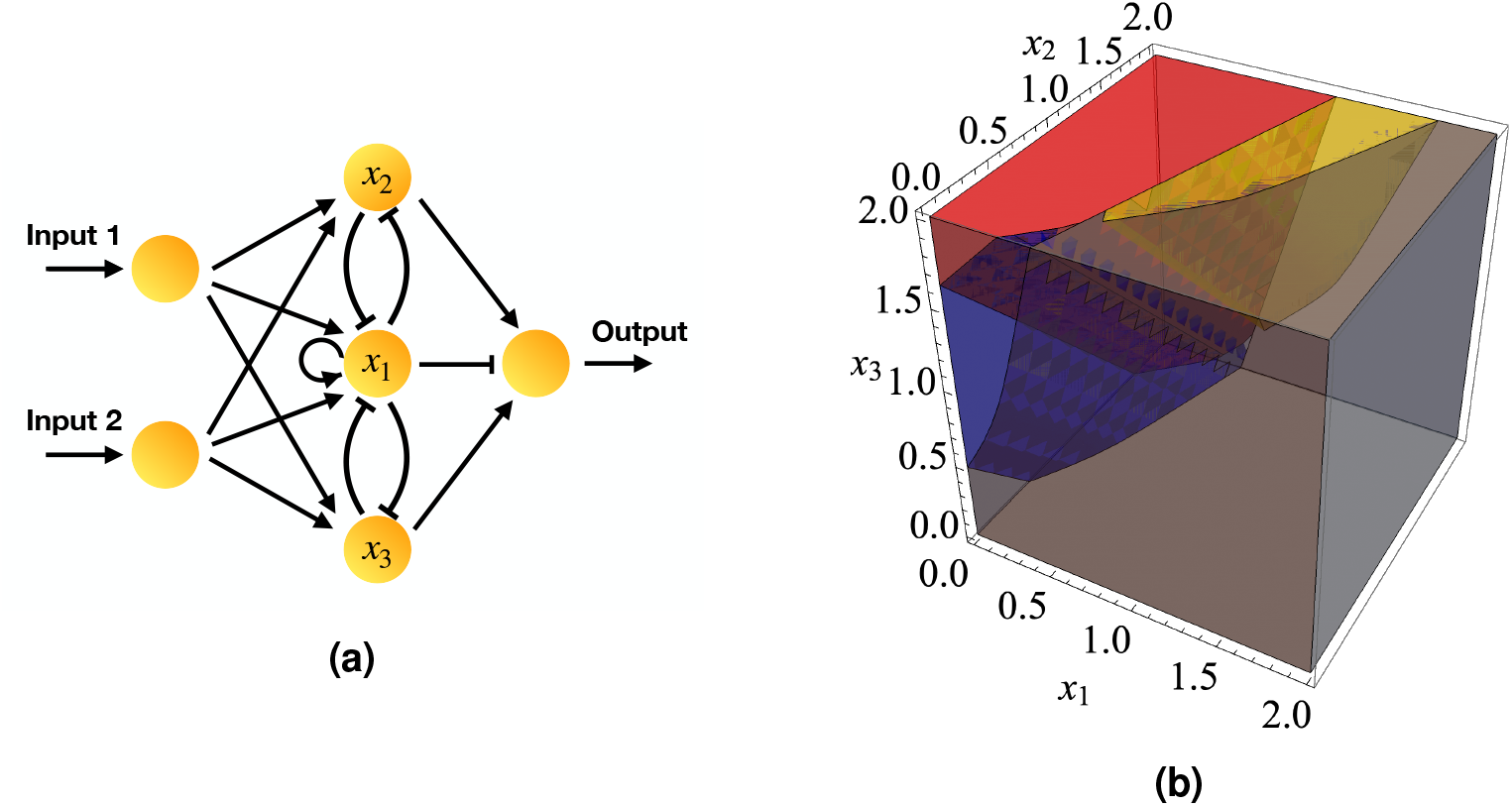
The XORANDOROFF gate. **(a)** A multistable circuit which can dynamically switch among XOR, AND, OR and OFF logical functions. **(b)** The regions of phase space corresponding to each function. The colors yellow, red, gray, and blue correspond to the function AND, OR, OFF, and XOR, respectively.

### 2.4 Switchable binary adder/subtractor: an example

Let us demonstrate the applicability and power of the proposed framework for reducing the number of required elements by designing a circuit capable of performing binary addition as well as binary subtraction^28^. The traditional static circuits for either of these functions are depicted in Fig. 8a and 8b.

**Figure 8.**
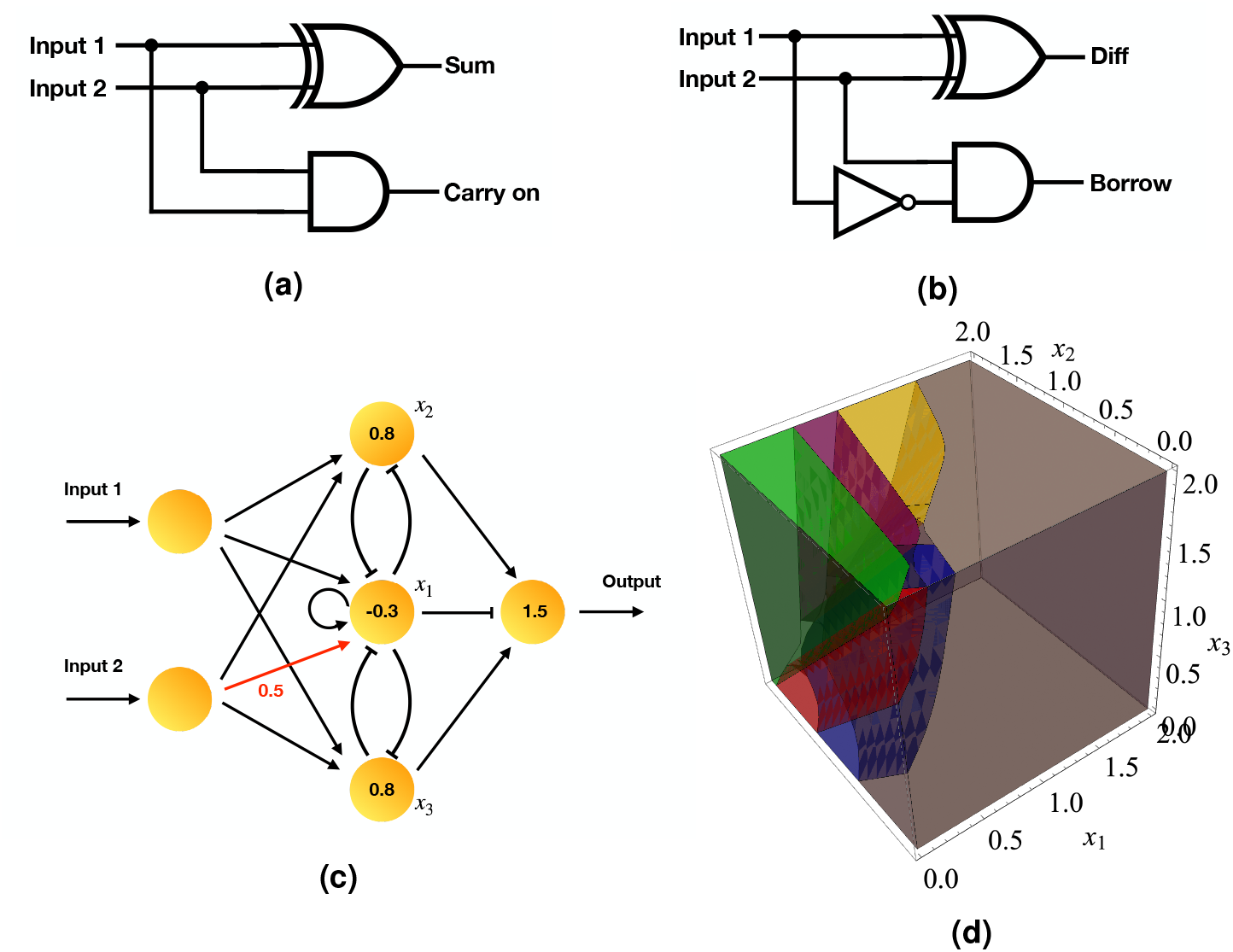
Traditional binary **(a)** adder and **(b)** subtractor circuits based on static components. **(c)** Dynamically switchable binary adder/subtractor circuit. **(d)** Initiating the system from each region shown here, enables it to perform a distinct function. Starting from green results in OR function, from red results in XOR, from blue performs NOT1+AND, from yellow does AND and if starting from grey the output is always OFF. Finally, starting from the purple area, the output simply reflects Input2.

As one can see in Fig. 8a, the adder circuit requires XOR and AND gates. Moreover, the binary subtractor shown in Fig. 8b needs XOR, NOT and AND. Being able to perform these two functions (i.e. addition and subtraction) in a traditional static framework, requires having these circuits in the system and rerouting signal (e.g. using other controller circuits) to the desired one when needed. However, if we allow context-dependency and reusing of the components, only three logic functions are required: AND, XOR and NOT1+AND. In the previous section, we already showed that having three nodes in the intermediate layer enables the system to perform AND, XOR and more. By breaking the symmetry of the inputs to the intermediate layer and adjusting the other parameters, one may modify this circuit into the one shown in Fig. 8c which is capable of performing the three logic functions required for binary adder/subtractor. This circuit is also able to perform the OR function, act as an OFF switch or just simply reflect input 2 as the output.Similar to the ANDOR gate, discussed in Sec. 2.2.1, switching among different functions in this circuit can be achieved by various means depending on the problem at hand.

Using the proposed circuit reduces the number of required elements and it’s therefore more efficient in this sense. However, it may sacrifice the speed of computation since there’s only one output from the system. This means that only either Sum or Carry (in binary adder) can be computed at a time and in the subtractor circuit, either Difference or Borrow. In order to circumvent this drawback one may alter the circuit structure and some parameters to get both outputs together, but this is beyond the scope of our study here, since we focus on maximum multifunctionality and reusibility of the elements.

## 3 Discussion

We here have designed and demonstrated three examples of dynamically switchable logic gates, based on multi-stability in the dynamics of regulation of signaling biochemical species, which switch between two, three and four distinct functions. We investigated the parameter ranges in which the conditions for this multi-functionality are met. Using the theory of large deviations, we characterized noise induced transitions between the possible logical functions and determined the reliability of the decisions. We therefore demonstrated that the proposed dynamical logic gates are resilient against three types of uncertainty: uncertainty in initial conditions, in control parameters, and in dynamics. Finally, we provided an example in which different switchable logic units can be combined to create a more complex computation which can be dynamically switched to a different, similarly complex computation.

We believe that these are only the first steps towards a more complete and faithful understanding of how network topology, dynamical systems, and information processing can combine in powerful, flexible, and non-trivial ways in the context of gene regulatory networks, signaling networks, and chemical computations. Many questions and implications remain to be explored. Thus far we have considered these biochemical dynamics playing out under the assumption of a well-mixed reaction volume, but spatial localisation and compartmentalisation are more and more appreciated as being important players in cell biology^29^,30 and could have significant effects on the operation of these switchable information processing elements. Another important direction will be to consider how families of more complex calculations built from the ANDOR gate and its cousins can be coupled to adaptive pressures and response on evolutionary timescales. It certainly seems plausible that the restrictions on total chemical signaling species number could make building different computations out of the same elements, as would be possible here, an attractive possibility. Finally, of course, the search is on for in vivo examples of ANDOR gates or higher order switchable computation.

Beyond the direct follow up questions, though, deeper implications also present themselves. Could one take advantage of these switchable gates to build an analog, chemical computer version of a deep learning network? Might biology have already done something similar? One might even imagine that chemical computations made flexibly switchable by the simple but non-trivial combination of network topology and reaction dynamics we propose here could have played a key role in the initiation of adaptation and natural selection at the origin of life.

## 4 Methods

### 4.1 Cellular regulatory dynamics

We here review general regulatory dynamics in biological systems. The time evolution of the concentration of the expressed gene *x* is governed by a simple, nonlinear dynamical equation composed of two terms corresponding to the production *F* (*s*) and degradation *λx*, i.e.

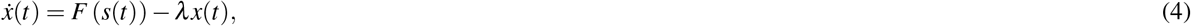

when the transcription process is faster than translation^31^. Here, *λ* is the degradation rate, *s* is the sum of all incoming regulatory signals (inductive, inhibitory, and self-regulation loops) to this gene, and *F* (*s*) is the regulatory function that describes the response of the expression rate to the input regulatory signal *s*. Throughout this study, we use a phenomenological and commonly used sigmoidal function

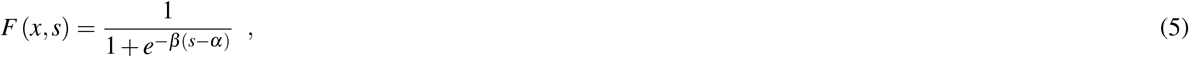

where *β* controls the sharpness (i.e. inverse of the fuzziness), and *α* controls the location of sigmoid function or in other words, the threshold in the signal after which the gene will be expressed^15^. It should be noted that the threshold *α* can also acquire negative values which means that the gene will be expressed even if it is inhibited with a strength smaller than this negative value.

Although we are using a specific type of dynamics for our examples which is shown in Eq. 5, one can in principle use any other type of regulatory function (e.g. the Hill function) as long as it features a switch-like sigmodal behavior and dynamically switching should still be achievable. In order to demonstrate this, we also constructed another version of the ANDOR gate based on the Hill function regulatory dynamics instead of the one shown in Eqs. 1 and 2:

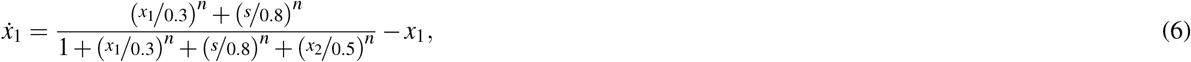

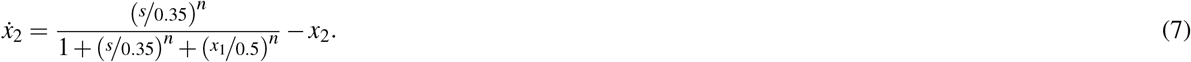

Assuming this dynamics for the intermediate layer of the ANDOR gate with *n* = 15 results in a phase portrait that is qualitativey similar to that of Eqs. 1 and 2. In Fig. 9, one can see these two plots side by side for the intermediate input signal level *s* = 1.

**Figure 9.**
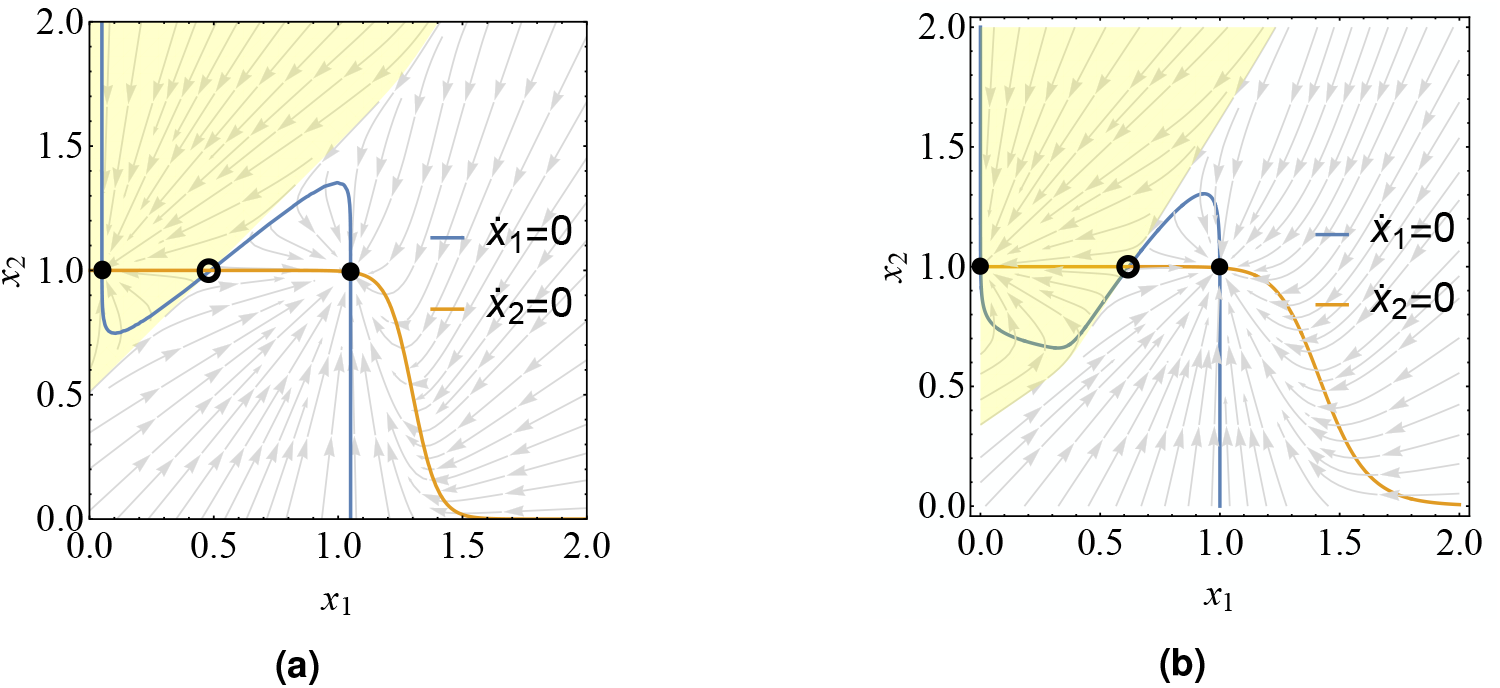
The phase portraits of the bistable motif in Fig. 2 **(a)** with the phenomenological dynamics and in **(b)** with Hill function dynamics. In both of these plots, the solid black circles show the fixed points, the hollow ones show the saddle points and the shaded area show the basins of attraction of the left fixed points.

### 4.2 Noise induced transitions in dynamical systems

One can define the dynamics of a stochastic system by a Langevin equation, i.e.

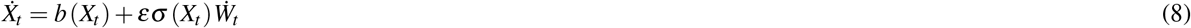

where the *n*-dimensional vector *X*_*t*_ is the state variable at time *t, b* (*X*_*t*_) is the deterministic part of the dynamics (i.e. drift vector), and the vector *W*_*t*_ is a *m*-dimensional Wiener process. Moreover, *ε* is a small parameter determining the noise strength, and *σ* (*X*_*t*_) is an *n × m* matrix, known as the diffusion matrix which determines the standard deviation of the noises and their contributions to each component of the dynamics. For such systems, the Freidlin-Wentzell action can be written as

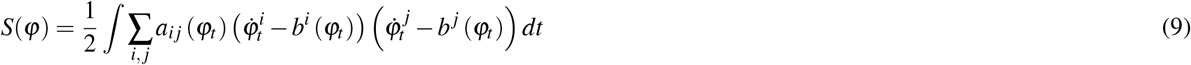

where (*a*_*i j*_(*x*)) = (*σ* (*x*)*σ*^*∗*^(*x*))^*−*1^. This action determines the difficulty of taking the path *φ*_*t*_ by the system. The probability for this path to be taken at a given noise strength *ε* is proportional to 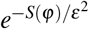. Obviously, for small noise strengths *ε*, all paths have negligible probability compared to the one which minimizes action *S*(*φ*).

Minimizing the action in Eq. 9 over the function space containing all paths which connect the point *X*_0_ at time *t* = 0 to *X*_*T*_ at time *t* = *T* provides the Minimum Action Path (MAP), and then, the minimum action value can be used for calculation of the rate of that transition. In order to find the transition probability from one state to another, one may intuitively expect to do this procedure for the transitions from every point in one basin of attraction to the fixed point of the other basin. However, it has been shown that all trajectories which leave a basin of attraction due to the fluctuations will visit a small neighborhood around the fixed point before leaving the basin^32^. Therefore, in the case of small fluctuations, the transition from one fixed point to the other represents the dominant transition and suffices for determining the resilience (against noise) of approaching the desired fixed point when starting from any point in its basin of attraction. We thus use this measure for studying the reliability of the decisions of our gate. It should be noted that when the system undergoes a transition from one fixed point to another, it spends most of the time at the fixed points and a small fraction of total time will be spent on the actual transition. Therefore, in order to get an acceptable accuracy, one needs to use an adaptive minimum action path method in which the distance (i.e. meshing) of time points is adaptively adjusted based on the speed^33^.

## Author contributions statement

This study is designed and conducted by M.B. and C.M.. The numerical work is done by M.B. and the manuscript is written and revised by M.B and C.M.

